# Autistic traits and suicidal thoughts, plans and self-harm in late adolescence: population based cohort study

**DOI:** 10.1101/165860

**Authors:** Iryna Culpin, Becky Mars, Rebecca M. Pearson, Jean Golding, Jon Heron, Isidora Bubak, Peter Carpenter, Cecilia Magnusson, David Gunnell, Dheeraj Rai

## Abstract

**Importance:** There have been recent concerns about a higher incidence of mortality by suicide in people with **a**utism spectrum disorder (ASD). To our knowledge, no large cohort studies have examined which features of autism may lead to suicidal ideation and behaviour, and whether there are any potential modifiable mechanisms.

**Objective:** To examine the hypothesis that ASD diagnosis and traits in childhood are associated with suicidal thoughts, plans and self-harm at 16 years, and that any of the observed associations are explained by depression in adolescence at 12 years.

**Design, setting and participants:** Prospective investigation of associations between ASD diagnosis and autistic traits with suicidal ideation and behaviour and a potential risk pathway via depression in early adolescence in 5,031 members of the Avon Longitudinal Study of Parents and Children.

**Main outcomes and measures:** History of self-harm with and without suicidal intent, suicidal thoughts and plans at 16 years assessed using a detailed self-report questionnaire. Exposures were ASD diagnosis and four measures (the coherence subscale of the Children’s Communication Checklist, the Social and Communication Disorders Checklist, a repetitive behaviour measure, and the sociability temperament subscale of the Emotionality, Activity and Sociability scale) dichotomised to represent the autism trait groups. Depressive symptoms in early adolescence were measured by the Short Moods and Feelings Questionnaire at 12 years.

**Results:** Children with impaired social communication had a higher risk of self-harm with suicidal intent (RR 2.10, 95% CI 1.28, 3.34), suicidal thoughts (1.42 times (95% CI 1.06, 1.91) and suicidal plans (RR 1.95, 95% CI 1.09, 3.47) by the age of 16 years as compared to those without. There was no evidence for an association between ASD diagnosis and the outcomes although these analyses were imprecise due to small numbers. There was also no evidence of an association between other autism trait measures and the outcomes. Approximately 32% of the total estimated association between social communication impairment and self-harm was explained by depressive symptoms at age 12 years.

**Conclusions:** Impairments in social communication are important in relation to suicidality. Early identification and management of depression may be a preventative mechanism and future research identifying other modifiable mechanisms may lead to preventative action or interventions against suicidal behaviour in this high-risk group.

## Introduction

Autism Spectrum Disorders (ASD) are developmental disorders characterised by deficits in social interaction and communication, restricted range of interests and repetitive behaviours.^1^ An increase in premature mortality in this population has been recently reported, with suicide being suggested as a significant contributor.^2^ However, there is a lack of population-based research on suicidal behaviour and suicidal ideation in this population.^3^ Suicidal behaviour describes self-harm (with or without suicidal intent) and completed suicide,^4^ whilst suicidal ideation refers to suicidal thoughts and cognitions.^3^ Self-harm and suicidal behaviour are highly prevalent in young people,^5,6^ and self-harming behaviours are a strong risk factor for completed suicide.^7^

The possibility of higher rates of suicidal ideation and attempts in individuals with ASD has been reported,^8,9^ however the existing research has mostly focused on either case reports^10^ or cross-sectional studies carried out in clinical^11, 12^ and non-clinical settings.^13^ The cross-sectional design,^11-13^ selective nature of the samples,^11^ and lack of adequate comparison group^12^ increase the likelihood of selection bias and limits generalisability of these findings. There remains a lack of epidemiological research using large population-based samples whilst accounting for possible confounding factors. It is also important to distinguish between self-harm with and without suicidal intent as these, although related, are clinically distinct outcomes.^14^ In addition, a growing body of research argues that the social, communication and behavioural difficulties comprising the autism spectrum may have distinct aetiologies,^15^ and it is plausible that outcomes related to difficulties in individual autistic traits may also differ. To our knowledge, there have been no prospective cohort studies examining the association between autistic traits and suicidal behaviour and ideation.

Furthermore, the mechanisms underlying any associations between autism/autistic traits and suicidality have not been examined. For example, depression is a strong risk factor for suicidal ideation^16^ and self-harm^17^ in the general population, however whether it could explain a greater risk of suicidal thoughts or behaviours in people with autism has not been studied. Quantifying this relationship is important since it may inform preventative or intervention strategies considering depression is potentially treatable. We used prospectively collected data from the Avon Longitudinal Study of Parents and Children (ALSPAC), a large birth cohort in Bristol, to address some of these gaps in the literature. Our research questions were:

1. Is an autism diagnosis and/or autistic traits associated with suicidal ideation (suicidal thoughts and plans) and suicidal behaviour (self-harm with and without suicidal intent) by age 16 years?
2. Are any of the observed associations explained by depressive symptoms in early adolescence?

## Methods

### Data source

The sample comprised participants from the Avon Longitudinal Study of Parents and Children (ALSPAC). During Phase I enrolment, 14,541 pregnant mothers residing in the former Avon Health Authority in the south-west of England with expected dates of delivery between 1 April 1991 and 31 December 1992 were recruited. These pregnancies resulted in 14,062 live births and 13,988 were alive at 1 year of age. When the oldest children were approximately 7 years of age, an attempt was made to bolster the initial sample with eligible cases who had failed to join the study originally. The total sample size for analyses using data after the age of seven is 15,247 pregnancies, of which 14,701 were alive at 1 year of age. Detailed information about the cohort has been collected since early pregnancy, including regular self-completion questionnaires from mothers and children. Information about ALSPAC is available at www.bristol.ac.uk/alspac/, including a searchable data dictionary (http://www.bris.ac.uk/alspac/researchers/data-access/data-dictionary/). For further details on the cohort profile, representativeness and phases of recruitment see Boyd et al.^18^

### Measures

#### Autism Spectrum Disorders: ASD diagnosis and autistic spectrum traits

Identification of children with diagnosed ASD in ALSPAC has been described elsewhere.^19-21^ A multisource approach included a record linkage study identifying cases from (i) community paediatric records; (ii) autism as the main reason for special educational needs from school records; (iii) maternal reports at age 9 years that the child had been diagnosed with ‘an autistic spectrum disorder or Asperger syndrome’; (iv) free text questionnaire responses from 6 months to 11 years; (v) ad hoc letters from parents to the Study Director.^19^ The diagnosis of ASD in ALSPAC has been previously validated by a consultant paediatrician using the International Classification of Diseases (ICD-10),^20^ and we have cross validated cases ascertained from maternal reports against autistic spectrum traits.^21^

Four individual measures optimally predictive of autism diagnosis in ALSPAC^22^ via parental questionnaires were analysed. These included the Social and Communication Disorder Checklist (91 months),^23^ a measure of repetitive behaviour (69 months),^24^ the sociability subscale of the Emotionality, Activity and Sociability (38 months) temperament scale,^25^ and the coherence subscale of the Children’s Communication Checklist (115 months).^26^ Each ASD trait was dichotomised at approximately 10% cut-off to identify the high-risk group as described elsewhere.^21^

#### Self-harm, suicidal thoughts and plans

*S*elf-harm questions were based on those used in the Child and Adolescent Self-Harm in Europe Study (CASE).^5^ Participants who responded positively to the question ‘Have you ever hurt yourself on purpose in any way (e.g., by taking an overdose of pills or by cutting yourself?)’ at 16 years were classified as having a lifetime history of self-harm. Responses to two additional questions were used to identify those who self-harmed with suicidal intent: (i) selecting the response option ‘I wanted to die’ in response to the question ‘Do any of the following reasons help to explain why you hurt yourself on that (i.e. the most recent), occasion? or (ii) a positive response to the question ‘On any of the occasions when you have hurt yourself on purpose, have you ever seriously wanted to kill yourself? These questions enabled to identify individuals who had harmed with suicidal intent at some point during their lifetime, and those who had only ever engaged in non-suicidal self-harm. Self-harm behaviours were classified according to individual’s self-reported suicidal intent.^27^

Lifetime history of suicidal thoughts and plans were also assessed with the following questions at 16 years: ‘Have you ever thought of killing yourself, even if you would not really do it?’ and ‘Have you ever made plans to kill yourself?’.

#### Mediating variable

To examine whether depressive symptoms in early adolescence mediate the association between childhood ASD and suicidal behaviour in late adolescence we used data from the Short Mood and Feelings Questionnaire (SMFQ), a 13-item instrument used to evaluate core depressive symptomatology in children aged 8 to 18 years,^28^ assessed at 12 years. The SMFQ correlates highly with more extensive depression rating scales and diagnostic tools such as the Children’s Depression Inventory^29^ and the Diagnostic Interview Schedule for Children.^30^

#### Confounding variables

Parental, socioeconomic, and family characteristics identified in previous studies as being associated with ASD and suicidal behaviour were collected prospectively from maternal questionnaires during the antenatal period. These included financial problems (occurrence of major financial problems since pregnancy *versus* none), highest maternal educational attainment (less than O-Level, O-Level, A-Level *versus* university degree), parental social class (professional/managerial *versus* manual) with the highest of maternal or paternal social class used; and accommodation type (detached house, semi-detached house *versus* flat); maternal age (in years), maternal antenatal (18 and 32 weeks’ gestation) and early postnatal (8 weeks and 8 months) depression assessed using the Edinburgh Postnatal Depression Scale (EPDS)^31^, maternal antenatal anxiety (18 and 32 weeks’ gestation) measured using anxiety items from the Crown-Crisp Index, a validated self-rating inventory;^32^ parental suicide attempt assessed using maternal questionnaires repeated eight times from birth to 11 years (yes or no); and sexual abuse (repeated seven times from birth to eight years) and physical cruelty (repeated eight times from birth to 11 years) to children in the household by mother/partner (yes or no). Analyses were also adjusted for child’s gender (male *versus* female) and ethnicity (white *versus* non-white).

### Statistical analyses

#### Main effects

First, we compared characteristics of missing children with those who comprised the study sample and carried out descriptive analysis. In the main analysis, we used multinomial regression to examine associations with explanatory variables and a three-category self-harm outcome: no self-harm; self-harm without suicidal intent; and self-harm with suicidal intent at 16 years. We used modified Poisson regression to examine the associations between explanatory variables and suicidal thoughts and plans at 16 years as binary outcomes to derive relative risks and robust 95% confidence intervals. A modified Poisson regression approach (with a robust error variance) directly estimates relative risks and robust error estimates with binary outcomes.^33^ All missing data were imputed, and all analyses were repeated using data for n=5,093 (main effects) and n=7,788 (mediation) adolescents. We imputed for missing data because ignoring those with missing data can result in bias by making the assumption that data are missing completely at random.^34^ A complete description of the imputation method is presented in the eMethod in the Supplement. We tested models unadjusted and adjusted for the potential confounding factors. In accordance with ALSPAC policy to protect confidentiality, analyses wherein the number of participants was less than 5 in a given cell were censored. All analyses were conducted using Stata v.13 (StataCorp., USA).

#### Mediating effects

We wanted to assess the importance of depressive symptoms at age 12 years in explaining the association between autistic traits in childhood and self-harm at age 16 years. Direct (ASD to risk of self-harm) and indirect (through depressive symptoms) pathways were estimated using Structural Equation Modelling (SEM) in Mplus v.7.^35^ Confirmatory Factor Analysis (CFA) was used to derive a normally distributed latent trait underlying the observed SMFQ^28^ scores using ordinal response items. A latent trait approach helps account for measurement error and increases power by modelling variables as a continuous trait.^36^ The approach recommended by Muthén^37^ was used to estimate mediation effects within the context of possible confounding. Analyses were adjusted for a range of individual, maternal and familial confounders. Detailed description of the mediation method is presented in the Supplement.

## Results

### Sample derivation

Our starting sample included those with data on at least one ASD exposure (presence/absence of ASD diagnosis or data on at least one of the four dichotomised ASD traits; n=14,684). Complete outcome data on suicidal behaviour and ideation were available for 5,031 adolescents. A description of the numbers of participants available with data on suicidal behaviour and ideation, ASD exposures and confounders is shown in Figure 1. The descriptive statistics of the sample by the presence of autism/autistic traits are presented in eTable1. Characteristics of the sample by the completeness of data availability are presented in eTable2.

**Figure 1.**
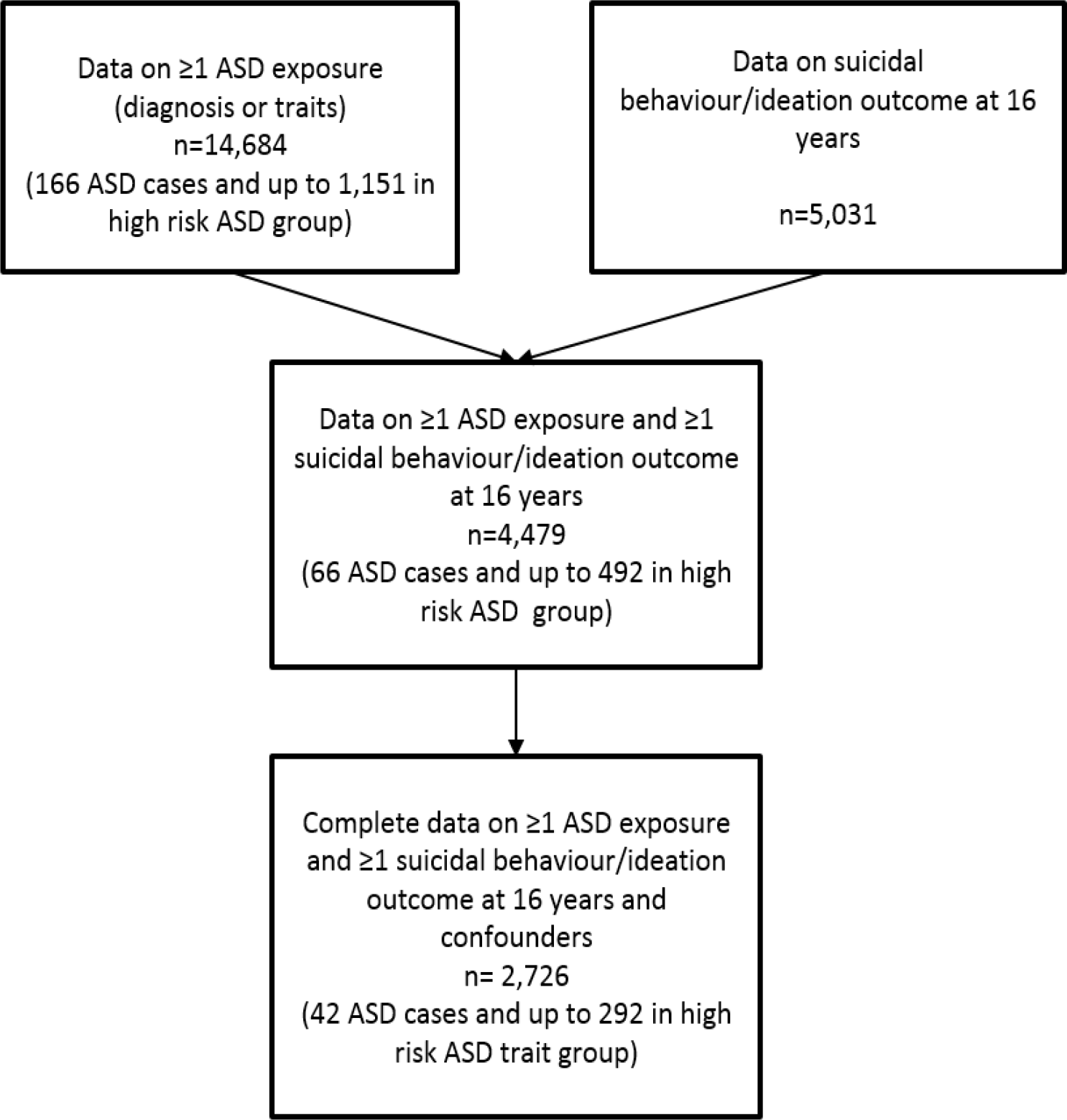
Study sample. Note: ASD=Autism Spectrum Disorder.

**Table 1.** Prevalence of self-harm with and without suicidal intent at 16 years in young adults by Autism Spectrum Disorder and autistic trait measures

| Exposure/Risk group<br>(age of assessment) | No self-harm<br>(n=4,114) | Self-harm<br>without<br>suicidal<br>intent<br>(n=602) | Self-harm<br>with suicidal<br>intent<br>(n=347) | P-value |
| --- | --- | --- | --- | --- |
| ASD diagnosis (diagnosed by 11 years),<br>n (%) |  |  |  | - |
| No ASD diagnosis | 4,054 (81.2%) | 595 (11.9%) | 343 (6.9%) |  |
| ASD diagnosis | 60 (84.5%) | Censored | Censored |  |
| Reduced social cognition (91 months),<br>n (%) |  |  |  | <0.001 |
| Lower risk | 3,220 (82.3%) | 460 (11.8%) | 233 (5.9%) |  |
| Higher risk | 249 (76.2%) | 37 (11.3%) | 41 (12.5%) |  |
| Repetitive behaviour (69 months),<br>n (%) |  |  |  | 0.074 |
| Lower risk | 3,254 (81.6%) | 477 (11.9%) | 258 (6.5%) |  |
| Higher risk | 212 (80.3%) | 26 (9.8%) | 26 (9.9%) |  |
| Reduced sociability (38 months),<br>n (%) |  |  |  | 0.833 |
| Lower risk | 3,257 (81.7%) | 463 (11.6%) | 269 (6.7%) |  |
| Higher risk | 408 (82.8%) | 54 (10.9%) | 31 (6.3%) |  |
| Reduced coherence (115 months),<br>n (%) |  |  |  | <0.001 |
| Lower risk | 3,311 (81.7%) | 491 (12.1%) | 250 (6.2%) |  |
| Higher risk | 302 (79.7%) | 34 (9.0%) | 43 (11.3%) |  |

*Note:* p-values based on Chi-Square test of the association between self-harm with/without suicidal intent and categorical ASD exposures.
Censored to prevent disclosure due to small cell counts.

**Table 2.**
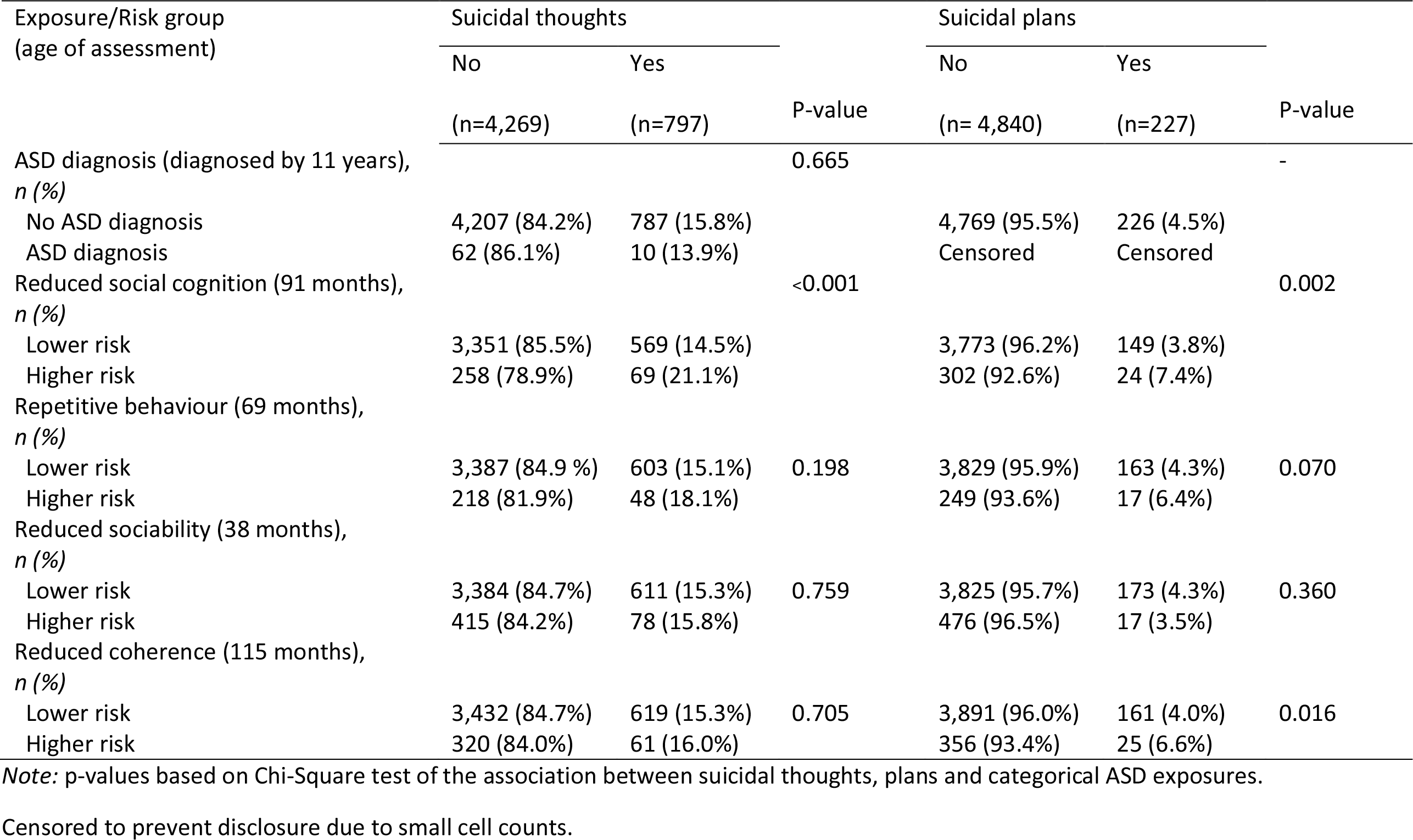
Prevalence of suicidal thoughts and plans at 16 years in young adults with Autism Spectrum Disorder and autistic trait measures

| Exposure/Risk group<br>(age of assessment) | Suicidal thoughts |  |  | Suicidal plans |  |  |
| --- | --- | --- | --- | --- | --- | --- |
|  | No | Yes | P-value | No | Yes | P-value |
|  | (n=4,269) | (n=797) |  | (n= 4,840) | (n=227) |  |
| ASD diagnosis (diagnosed by 11 years),<br>n (%) |  |  | 0.665 |  |  | - |
| No ASD diagnosis | 4,207 (84.2%) | 787 (15.8%) |  | 4,769 (95.5%) | 226 (4.5%) |  |
| ASD diagnosis | 62 (86.1%) | 10 (13.9%) |  | Censored | Censored |  |
| Reduced social cognition (91 months),<br>n (%) |  |  | <0.001 |  |  | 0.002 |
| Lower risk | 3,351 (85.5%) | 569 (14.5%) |  | 3,773 (96.2%) | 149 (3.8%) |  |
| Higher risk | 258 (78.9%) | 69 (21.1%) |  | 302 (92.6%) | 24 (7.4%) |  |
| Repetitive behaviour (69 months),<br>n (%) |  |  |  |  |  |  |
| Lower risk | 3,387 (84.9 %) | 603 (15.1%) | 0.198 | 3,829 (95.9%) | 163 (4.3%) | 0.070 |
| Higher risk | 218 (81.9%) | 48 (18.1%) |  | 249 (93.6%) | 17 (6.4%) |  |
| Reduced sociability (38 months),<br>n (%) |  |  |  |  |  |  |
| Lower risk | 3,384 (84.7%) | 611 (15.3%) | 0.759 | 3,825 (95.7%) | 173 (4.3%) | 0.360 |
| Higher risk | 415 (84.2%) | 78 (15.8%) |  | 476 (96.5%) | 17 (3.5%) |  |
| Reduced coherence (115 months),<br>n (%) |  |  |  |  |  |  |
| Lower risk | 3,432 (84.7%) | 619 (15.3%) | 0.705 | 3,891 (96.0%) | 161 (4.0%) | 0.016 |
| Higher risk | 320 (84.0%) | 61 (16.0%) |  | 356 (93.4%) | 25 (6.6%) |  |

Note: p-values based on Chi-Square test of the association between suicidal thoughts, plans and categorical ASD exposures.
Censored to prevent disclosure due to small cell counts.

### Main effects

Of 5,031 adolescents with data on suicidal behaviour and ideation up to age 16 years, 601 (11.9%, 95% CI 11.0 to 12.8%) reported self-harm without suicidal intent, 347 (6.9%, 95% CI 6.2% to 7.6%) reported self-harming with suicidal intent, 797 (15.7%, 95% CI 14.6 to 16.4) reported experiencing suicidal thoughts, and 227 (4.5%, 95% CI 3.8 to 5.0) reported making suicidal plans (Tables 1-2). The regression analysis (Table 3) provided evidence for an effect of impaired social communication on risk of self-harm with suicidal intent (adjusted RR 2.14, 95% CI 1.28 to 3.58), but not self-harm without suicidal intent (RR 1.02, 95%CI 0.62 to 1.67). There was also evidence for an effect of impaired social communication on risk of suicidal thoughts (RR 1.42, 95% CI 1.06 to 1.91) and suicidal plans (RR 1.95, 95%CI 1.09 to 3.47; Table 4). There was no evidence of an association between ASD diagnosis, although the numbers were very low and confidence intervals wide. None of the other autistic traits (sociability, coherence, and repetitive behaviour) appeared to be associated with the outcomes (Tables 3-4).

**Table 3.**
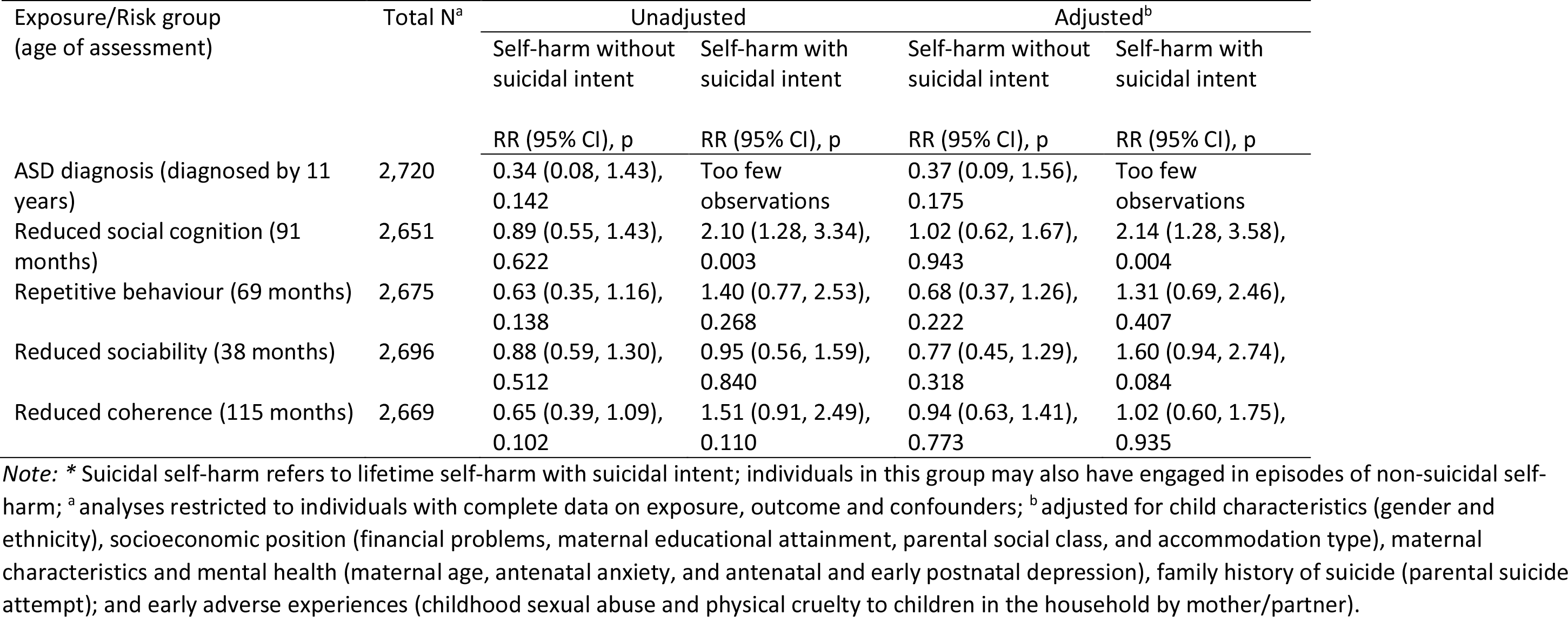
Associations between Autism Spectrum Disorder (ASD *versus* no ASD), autistic trait measures (high *versus* low ASD risk group) and self-harm with/without suicidal intent^*^ (*versus* no self-harm) at 16 years

**Table 4.** Associations between Autism Spectrum Disorder (ASD *versus* no ASD), autistic trait measures (high *versus* low ASD risk group) and suicidal thoughts (*versus* no suicidal thoughts) and plans (*versus* no suicidal plans) at 16 years

| Exposure/Risk group<br>(age of assessment) | Total N<sup>a</sup> | Unadjusted | Adjusted<sup>b</sup> | Total N<sup>a</sup> | Unadjusted | Adjusted<sup>b</sup> |
| --- | --- | --- | --- | --- | --- | --- |
|  |  | Suicidal thoughts |  |  | Suicidal plans |  |
|  |  | RR (95% CI), p |  |  | RR (95% CI), p |  |
| ASD diagnosis (diagnosed by 11 years) | 2,726 | 0.32 (0.08, 1.25),<br>0.101 | 0.43 (0.11, 1.73),<br>0.238 | 2,728 | Too few<br>observations | Too few<br>observations |
| Reduced social cognition (91 months) | 2,658 | 1.41 (1.04, 1.90),<br>0.024 | 1.42 (1.06, 1.91),<br>0.019 | 2,659 | 1.89 (1.07, 3.33),<br>0.027 | 1.95 (1.09, 3.47),<br>0.024 |
| Repetitive behaviour (69 months) | 2,681 | 1.23 (0.86, 1.76),<br>0.250 | 1.20 (0.85, 1.69),<br>0.299 | 2,683 | 1.41 (0.70, 2.87),<br>0.331 | 1.44 (0.75, 2.74),<br>0.270 |
| Reduced sociability (38 months) | 2,702 | 1.10 (0.83, 1.47),<br>0.493 | 1.16 (0.87, 1.55),<br>0.305 | 2,704 | 0.83 (0.42, 1.64),<br>0.601 | 0.87 (0.44, 1.72)<br>0.689 |
| Reduced coherence (115 months) | 2,674 | 0.84 (0.58, 1.22),<br>0.370 | 0.90 (0.63, 1.29),<br>0.577 | 2,676 | 1.0 (0.49, 2.03),<br>1.00 | 1.0 (0.49, 2.02),<br>0.979 |

*Note:* <sup>a</sup> analyses restricted to individuals with complete data on exposure, outcome and confounders; <sup>b</sup> adjusted for child characteristics (gender and ethnicity), socioeconomic position (financial problems, maternal educational attainment, parental social class, and accommodation type), maternal characteristics and mental health (maternal age, antenatal anxiety, and antenatal and early postnatal depression), family history of suicide (parental suicide attempt); and early adverse experiences (childhood sexual abuse and physical cruelty to children in the household by mother/partner).

Results were comparable when using imputed data sets for the main effect of impaired social communication on the outcomes (eTables 3-4). In these imputed analyses, there was evidence for the main effect of repetitive behaviour and reduced coherence on risk of self-harm with suicidal intent (RR 1.58, 95%CI 1.00 to 2.48 and RR 1.97 95%CI 1.36 to 2.81 respectively), and suicidal plans (RR 1.63, 95%CI 1.03 to 2.58 and 1.61, 95%CI 1.08 to 2.41 respectively).

### Mediation effects

Given the main effect of impaired social communication on risk of self-harm, we examined whether depressive symptoms in early adolescence mediate this association. In order to estimate the mediation model, we combined suicidal/non-suicidal self-harm into one category. First, we examined the fit of the measurement model incorporating exposure, mediator and confounders. The full model was run without using bootstrapping to enable calculation of model fit statistics. The Root Mean Square Error of Approximation (RMSEA=0.04, 95%CI 0.03 to 0.04) and the Comparative Fit Index (CFI=0.97) indicated that the measurement model fit the data well, supporting the adequacy of the model for tests of structural paths and mediation effects. Unadjusted and adjusted structural mediation models were estimated to examine the direct and indirect effect of impaired social cognition on self-harm through depressive symptoms whilst accounting for possible mediator-outcome confounding (Figure 2). There was evidence of an indirect pathway from impaired social cognition to self-harm via depressive symptoms (product coefficient [β]=0.087, 95% CI 0.03 to 0.14). There was little evidence of a direct pathway from impaired social cognition to self-harm once the indirect effect via depressive symptoms was accounted for (regression coefficient [β]=0.090, 95% CI −0.11 to 0.28). This indirect pathway via depressive symptoms accounted for 32% of the total estimated association between impaired social cognition and self-harm. Adjustment for the confounders made little difference to the parameter estimates (Table 5).

**Figure 2.**
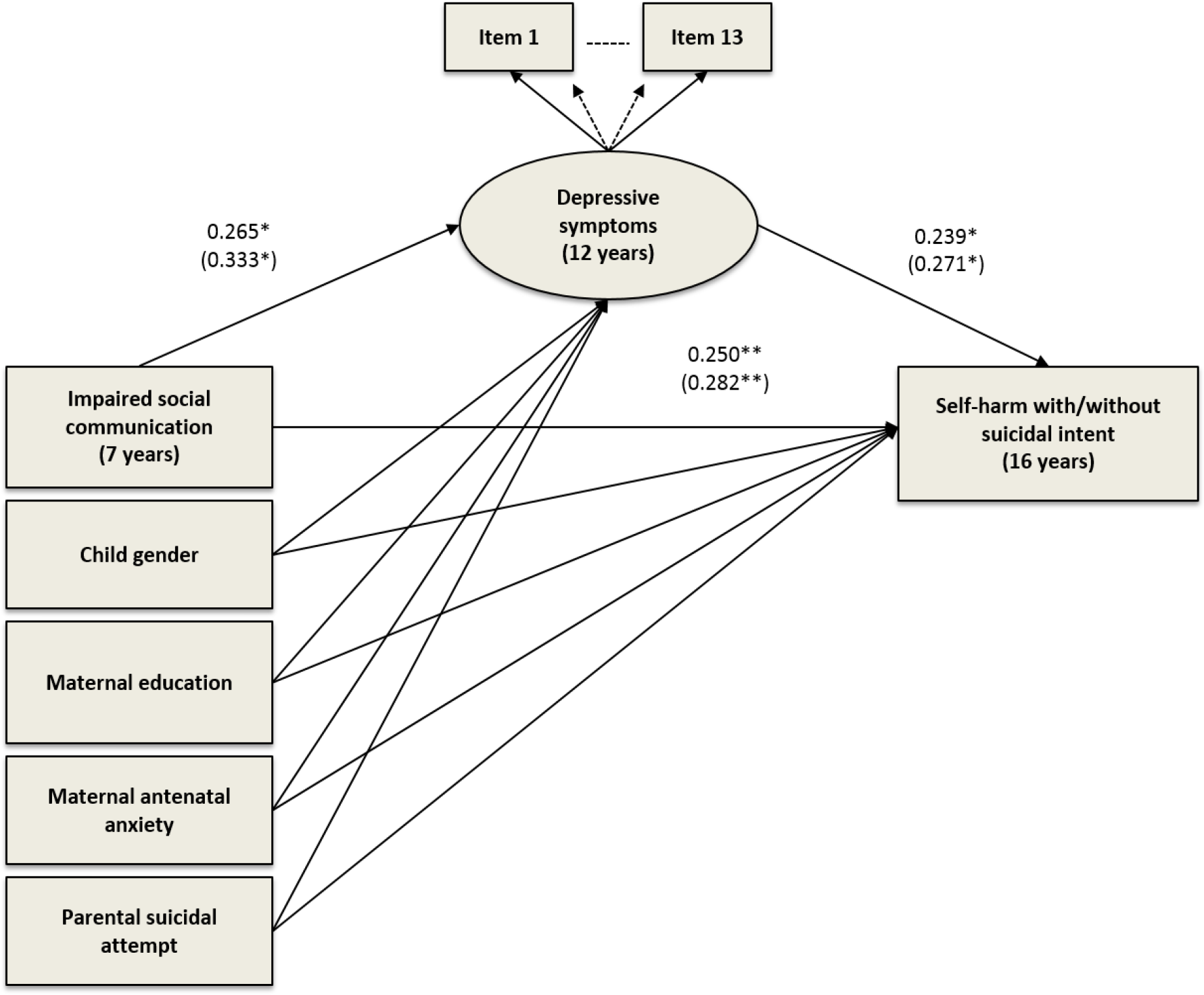
Structural mediation model estimating the direct effect of impaired social cognition on lifetime self-harm at 16 years and the indirect effect through child’s depressive symptoms at 12 years (adjusted for potential child, maternal and socioeconomic confounders). *Note*: observed variables are represented by squares, whilst the latent variable is represented by a circle. Covariances are not shown to reduce figure complexity. Paths coefficients in brackets are from the imputed data analysis.

^*^p≤0.001; and ^**^p<0.05

**Table 5.** Estimates of the direct effect and effect mediated through depressive symptoms in the association between impaired social cognition and self-harm at 16 years unadjusted and adjusted for antenatal confounders and child’s gender (n=2,936)^a^

| Effect Size <sup>b</sup> | Model estimates |  |  |  |
| --- | --- | --- | --- | --- |
|  | Unadjusted model |  | Adjusted model <sup>c</sup> |  |
|  | *B* (95% CI) | P Value | *B* (95% CI) | P Value |
| *1. Total effect* | 0.177 (-0.02, 0.38) | 0.084 | 0.250 (0.05, 0.45) | 0.016 |
| Low social cognition on self-harm |  |  |  |  |
| *2. Indirect effect* | 0.087 (0.03, 0.14) | 0.002 | 0.081 (0.03, 0.13) | 0.001 |
| Low social cognition on self-harm, through depressive symptoms |  |  |  |  |
| *3. Direct effect* | 0.090 (-0.11, 0.28) | 0.372 | 0.169 (-0.03, 0.37) | 0.102 |
| Low social cognition on self-harm, adjusted for depressive symptoms |  |  |  |  |

<sup>a</sup> Analyses restricted to participants with complete exposure, outcome, mediator and confounders
<sup>b</sup> *Effect Size*: Unadjusted and adjusted regression coefficients (unstandardised); *B*
<sup>c</sup> Analyses adjusted for child characteristics (gender), socioeconomic position (maternal educational attainment) and maternal characteristics (antenatal anxiety).

The direct and indirect estimates with imputed data sets led to similar results (eTable 5). However, the sizes of the observed direct and indirect effects were greater, which may suggest that attrition led to an underestimation of the direct and indirect effect sizes in the complete case analyses. Based on the imputed analyses, a slightly higher proportion (35%) of the total association between impaired social cognition and self-harm was accounted for by the indirect path through depressive symptoms.

## Discussion

To our knowledge, this is the first large population-based study to investigate the association between an autism diagnosis and traits and suicidal ideation and behaviour by late adolescence, as well as examining the mechanisms of this association. We did not find an association with diagnosed autism and the outcomes, although the estimates were imprecise due to small numbers (42 ASD cases). Impairments in social communication were associated with an increased risk of suicidal thoughts, suicidal plans and self-harm with suicidal intent, but not self-harm without suicidal intent.

Our findings suggest that social communication difficulties may play a central role in relation to suicidality and are consistent with existing case studies suggesting that social impairments and difficulties in establishing interpersonal relationships are triggers for suicidal behaviour.^10,38^ Suicidal behaviour in autistic individuals is often underreported,^39^ particularly in those with impaired communicative abilities and comorbid self-injurious behaviour.^40^ Our findings emphasise the potential importance of assessing whether self-injurious behaviour occurs in the context of suicidal ideation.^3^ The stronger associations for social communication with suicidality outcomes as compared to the other traits of autism is concordant with the argument on the fractionation of core autistic impairments.^15^

It has been argued that it is adolescents with ASD without intellectual disability who are at most risk of suicidal ideation and behaviour^41,42^ due to the increased awareness of their social difficulties and secondary depression associated with social isolation and exclusion.^11^ We were not able to directly test this possibility since participants in our sample were predominantly high-functioning individuals (96.4% of individuals with autism diagnosis had IQ>70) and we did not have enough statistical power to study lower- and higher-functioning individuals separately.

### Possible mechanism

We tested whether depression during early adolescence could explain the association between social communication and suicidal behaviour. Children with impaired social communication skills were at increased risk of depressive symptoms in early adolescence, which, in turn, was a strong risk factor for suicidal behaviour later in adolescence. It should be noted that although depression explained about a third of the variance of the association between childhood autistic traits and suicidal behaviour, substantial variance remained unexplained. This finding emphasises the need for identifying other potentially modifiable mechanisms in this relationship.

### Strengths and limitations

The strengths of this study include the large sample, the long-term follow-up, the availability of data on several outcomes, as well as rich data on confounders, and longitudinal design that enables to examine mediating pathways. Furthermore, we were able to examine a range of autistic traits in a large population,^3^ despite a relatively small number of cases with ASD diagnosis. The findings need to be interpreted in light of several limitations. Firstly, our study was likely to be underpowered to detect the association between ASD diagnosis and the outcomes due to a relatively small number of diagnosed cases. Secondly, despite the population-based sample, it is not possible to rule out selection bias in relation to baseline recruitment or attrition in the sample over time. We attempted to address this by controlling for factors known to be predictive of attrition in ALSPAC and by imputing missing data. The pattern of missing data and imputed analyses suggests that attrition may have led to an underestimation of the size of the association between ASD traits, in particular repetitive behaviour and impaired speech coherence, and suicidal ideation and behaviour. Thirdly, there are limitations in establishing suicidal intent accompanying self-harm, particularly using self-reports which could be influenced by fluctuations in mood or change over time. This could be further compounded in autistic individuals who may experience additional difficulties understanding or responding to such questionnaires.

In summary, this study suggests that children with impairments in social communication are at higher risk for suicidal ideation and behaviour in late adolescence. Depressive symptoms in early adolescence partially explain this association which emphasises the importance of addressing the mental health needs of autistic children. Future research is required to assess whether other modifiable mechanisms could be identified, as these may have the potential to lead to preventative action or interventions against suicidal behaviour in this high-risk group.

## Acknowledgements

This research was specifically funded by the Baily Thomas Foundation (Grant ref: 3747-6849). We are extremely grateful to all the families who took part in this study, the midwives for their help in recruiting them, and the whole ALSPAC team, which includes interviewers, computer and laboratory technicians, clerical workers, research scientists, volunteers, managers, receptionists and nurses. The UK Medical Research Council and Wellcome (Grant ref: 102215/2/13/2) and the University of Bristol provide core support for ALSPAC. This publication is the work of the authors who will serve as guarantors for the contents of this paper. This study was also supported by the NIHR Biomedical Research Centre at the University Hospitals Bristol NHS Foundation Trust and the University of Bristol. The views expressed in this publication are those of the author(s) and not necessarily those of the NHS, the National Institute for Health Research or the Department of Health. Dr Culpin is funded by the Elizabeth Blackwell Institute for Health Research Early Career Fellowship.

## Ethical approval

Ethical approval for the data collection was obtained from the ALSPAC Ethics and Law Committee and the Local Research Ethics Committees.

